# Ratio of the interferon-*γ* signature to the immunosuppression signature predicts anti-PD-1 therapy response in melanoma

**DOI:** 10.1101/2020.04.18.047852

**Authors:** Yan Kong, Canqiang Xu, Chuanliang Cui, Wenxian Yang, Shuang Yang, Zhihong Chi, Xinan Sheng, Lu Si, Yihong Xie, Jinyu Yu, Xuejun Chen, Shun Wang, Jing Hu, Frank Zheng, Wengang Zhou, Rongshan Yu, Jun Guo

**Affiliations:** Peking University Cancer Hospital and Institute, Beijing, China; Aginome Scientific Co., Ltd., Xiamen, China; Amoy Diagnostics Co., Ltd., Shanghai, China; XMU-Aginome Joint Lab, School of Informatics, Xiamen University, China

**Keywords:** Immunology, immunotherapy, anti-PD-1, biomarker, tumour microenvironment, immune checkpoint inhibitor

## Abstract

Immune checkpoint inhibitor (ICI) treatments produce clinical benefit in many patients. However, better pretreatment predictive biomarkers for ICI are still needed to help match individual patients to the treatment most likely to be of benefit. Existing gene expression profiling (GEP)-based biomarkers for ICI are primarily focused on measuring a T cell-inflamed tumour microenvironment that contributes positively to the response to ICI. Here, we identified an immunosuppression signature (IMS) through analysing RNA sequencing data from a combined discovery cohort (*n* = 120) consisting of three publicly available melanoma datasets. Using the ratio of an established IFN-*γ* signature and IMS led to consistently better prediction of the ICI therapy outcome compared to a collection of nine published GEP signatures from the literature on a newly generated internal validation cohort (*n* = 55) and three published datasets of metastatic melanoma treated with anti-PD-1 (*n* = 48) and anti-CTLA-4 (*n* = 42) as well as in patients with gastric cancer treated with antiPD-1 (*n* = 45), demonstrating the potential utility of IMS as a predictive/prognostic biomarker that complements existing GEP signatures for immunotherapy.

## 1 Introduction

Historically, advanced melanoma has a poor prognosis, with a 5-year survival rate of less than 10%^1^. Immune checkpoint inhibitors (ICIs) targeting PD-1 and CTLA-4 have shown improved survival in advanced melanoma patients^1–4^, but only a subset of patients respond. Additionally, the efficacy of ICIs has been observed to be significantly lower for East Asian melanoma patients than for Caucasian patients^5, 6^.

Published data suggest that tumour mutational burden (TMB) and PD-L1 expression may predict the clinical benefit of anti-PD-1 therapy in multiple cancer types^7–9^. Although the potential predictive power of PD-L1 expression for the clinical benefit of anti-PD-1 therapy remains controversial for advanced melanoma patients^10, 11^, higher TMB has been correlated with a superior clinical response^12, 13^, improved survival^14, 15^, and durable benefit^12, 16^ in advanced melanomas. In Asian melanoma patients, acral^17, 18^ and mucosal melanomas^18^ are the predominant subtypes and generally have a low point mutation burden. Consequently, it is not clear whether TMB is an effective predictor for advanced Asian melanoma patients.

In addition to TMB and PD-L1 expression, prediction models based on gene expression profiling (GEP) have also been proposed. Most GEP signatures consider T cell inflamed microenvironments characterized by the upregulation of IFN-*γ* signalling, antigen presentation, and immune checkpoint-related genes when predicting response to ICIs across multiple cancer types. However, these features are necessary but not always sufficient for a cancer patient to receive clinical benefit from ICI treatments^19^. A recent meta-review showed that predictive models built on inflamed GEP signatures achieved a moderate area under the receiver operator curve (AUC) value of 0.65 for the summary receiver operation characteristic (sROC) curve generated from 10 different solid tumour types in 8,135 patients^20^.

Here, we argued that immune suppressive elements in the tumour microenvironment (TME) should be considered in combination with an inflamed GEP signature to more accurately predict ICI therapy outcomes. The main objective of this study was to develop immunosuppressive GEP signatures that, when used in combination with inflamed GEP signatures, could better stratify patients based on their potential benefits from ICI therapy. We started by analysing RNA-seq data from baseline biopsy samples of melanoma patients treated with anti-PD-1 therapy and identified a set of 18 genes that played an “antagonistic” role against a pro-inflammatory TME and lead to negative outcomes in the discovery cohort consisting of multiple datasets. Our results reveal that key genes of the identified immunosuppression signature (IMS) are related to hallmark activities of cancer-associated fibroblasts (CAFs), macrophages and epithelial to mesenchymal transition (EMT), and the balance between the IFN-*γ* signature and the IMS plays an important prognostic and predictive role in both immunotherapy-naive primary tumours from The Cancer Genome Atlas (TCGA) database and ICI-treated patients.

## 2 Results

### Definition of an immunosuppression signature

We reviewed the literature and found three external datasets^14, 15, 21^ of advanced melanoma treated with an anti-PD-1 ICI with response and RNA-seq data, which we used as our discovery cohort (*n* = 120; Methods). We then identified 18 genes of which the expression levels, after adjusting for IFN-*γ* signature score, are consistently associated with negative response to ICI in the discovery cohort as our IMS (Fig. 1a-c). The genes in IMS with their respective biological functions are listed in Supplementary Table 1.

**Fig. 1.**
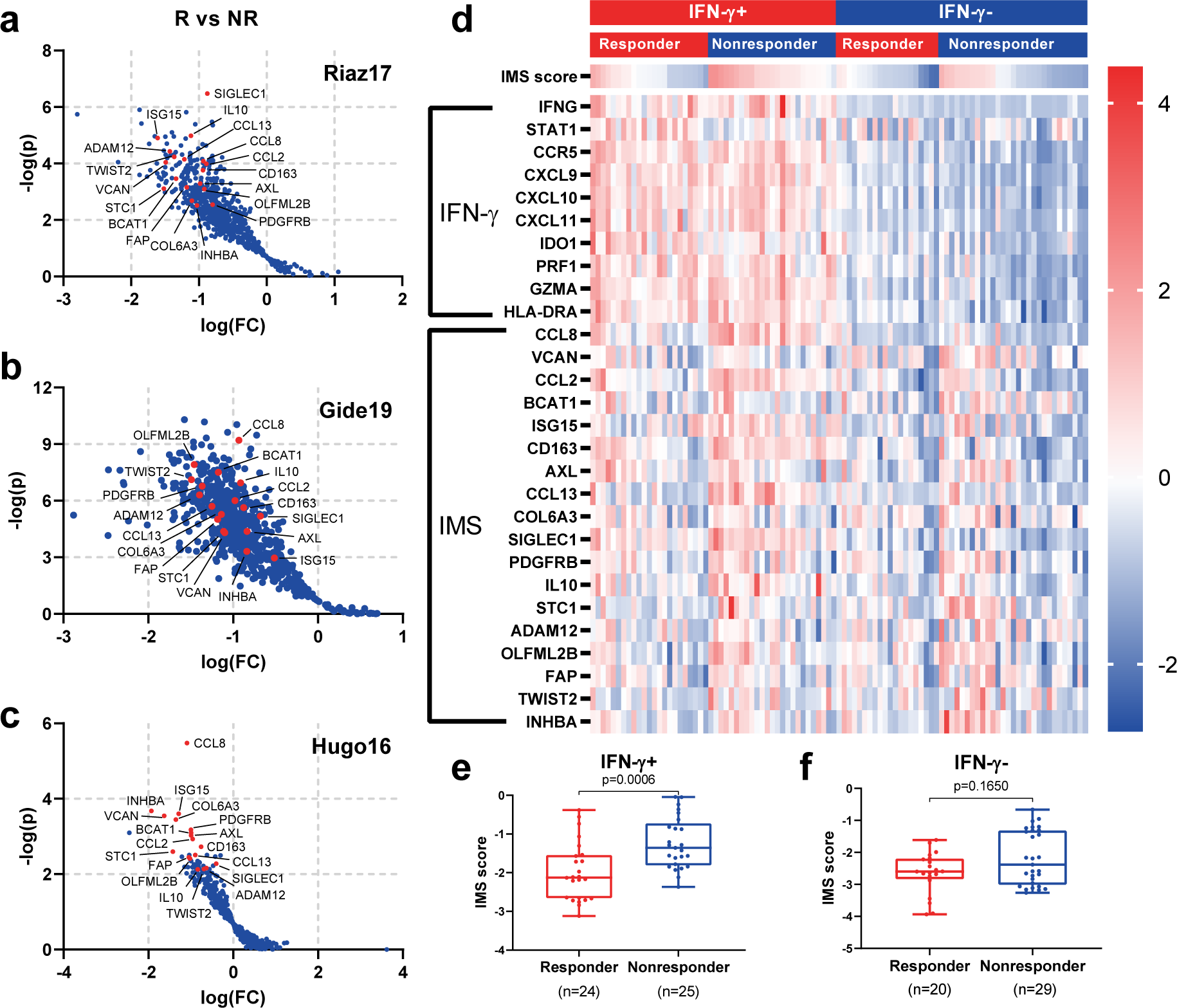
The definition of the IMS genes. **a-c**, Volcano plot depiction of differentially expressed genes after normalization by the IFN-*γ* score of individual sample by response on Riaz17 (in **a**, *n* = 51), Gide19 (in **b**, *n* = 41) and Hugo16 (in **c**, *n* = 28). R, responders (CR or PR); NR, nonresponders (PD) as per RECIST 1.1. IMS genes are highlighted in red. **d**, Heatmap showing the expression of genes from the IFN-*γ* signature and the IMS stratified by IFN-*γ* and response to anti-PD-1 therapy. Rows represent genes and columns represent patients. The expression levels were z-normalized within rows for visualization. The cutoff value for the IFN-*γ* signature score was set to its median. **e**, The IMS scores in responders versus nonresponders for IFN-*γ*+ and IFN-*γ*-subgroups.

Fig. 1d shows a heatmap of the expression of all genes in the IMS and IFN-*γ* signatures in the combined discovery cohort. Patients with elevated expression of IFN-*γ*-related genes included both responders and nonresponders, suggesting that an inflamed TME alone, as indicated by a higher IFN-*γ* signature score, is not sufficient to ensure positive outcomes from ICI. On the other hand, elevated expression levels of IMS genes were observed in patients from the nonresponder groups, particularly in the IFN-*γ*+ subgroup (*n* = 49, *p* = 0.0006). These data indicate a potential immunosuppressive role of the IMS genes that is opposite to the role of the IFN-*γ* related inflammatory signature, and both signatures should be considered in order to have more accurate predictions of outcomes from immunotherapy.

### Association of the immunosuppression signature with immune cell types

The identified IMS shows a strong presence of genes related to CAFs (FAP and PDGFRB^22^) and tumour-associated macrophages (TAMs) (CD163^23^ and SIGLEC1^24, 25^) as well as their associated cytokines or chemokines that lead to an immunosuppressive microenvironment (IL10^26^, CCL2, CCL8, and CCL13^14^) and stromal activities that lead to tumour proliferation, invasion and immune escape such as EMT or extracellular matrix (ECM) degradation (AXL^27^, TWIST2, ADAM12^28^, and COL6A3^29^). Therefore, high infiltration of CAFs and myeloid cells and their related stromal activities may be the reasons behind the lack of response from patients with an inflammatory TME. To further validate this hypothesis, we performed digital cell composition analysis using xCell^30^ on a combined melanoma dataset consisting of the three datasets in the discovery cohort and a TCGA melanoma dataset (*n* = 516), and analysed the distribution of different immune cell types with respect to the IFN-*γ* signature and IMS scores.

As expected, we observed that the IMS score was positively correlated with the abundances of fibroblasts (*r* = 0.62, *p* < 0.0001), monocytes (*r* = 0.45, *p* < 0.0001) and macrophages (*r* = 0.34, *p* < 0.0001) (Fig. 2a). Stratification of patients into IFN-*γ*+/- and IMS+/- subgroups according to their median values further revealed the different distributions of immune cells in relation to these two signatures (Fig. 2b). Fibroblasts were significantly enriched in IMS+ subgroups regardless of IFN-*γ* status (*p* < 0.0001; Fig. 2c). In addition, higher abundances of macrophage were associated with both higher IMS scores and higher IFN-*γ* signature scores. Interestingly, M2 macrophages, which play an important immunosuppressive role in the TME, were significantly associated with the IMS score in only the IFN-*γ*+ subgroups (*p* = 0.0281) but not the IFN-*γ*-subgroups. On the other hand, higher IFN-*γ* signature scores were associated with increased infiltration of CD8+ T cells, CD4+ T cells and B cells. However, the association of IMS scores and abundances of these cells within the microenvironment is not significant. All these results are consistent with the notion that IMS genes are related to immunosuppressive activities in cancers, and the balance between IFN-*γ* signature and IMS scores has a significant role in determining which patients benefit from adaptive immune rejuvating therapies.

**Fig. 2.**
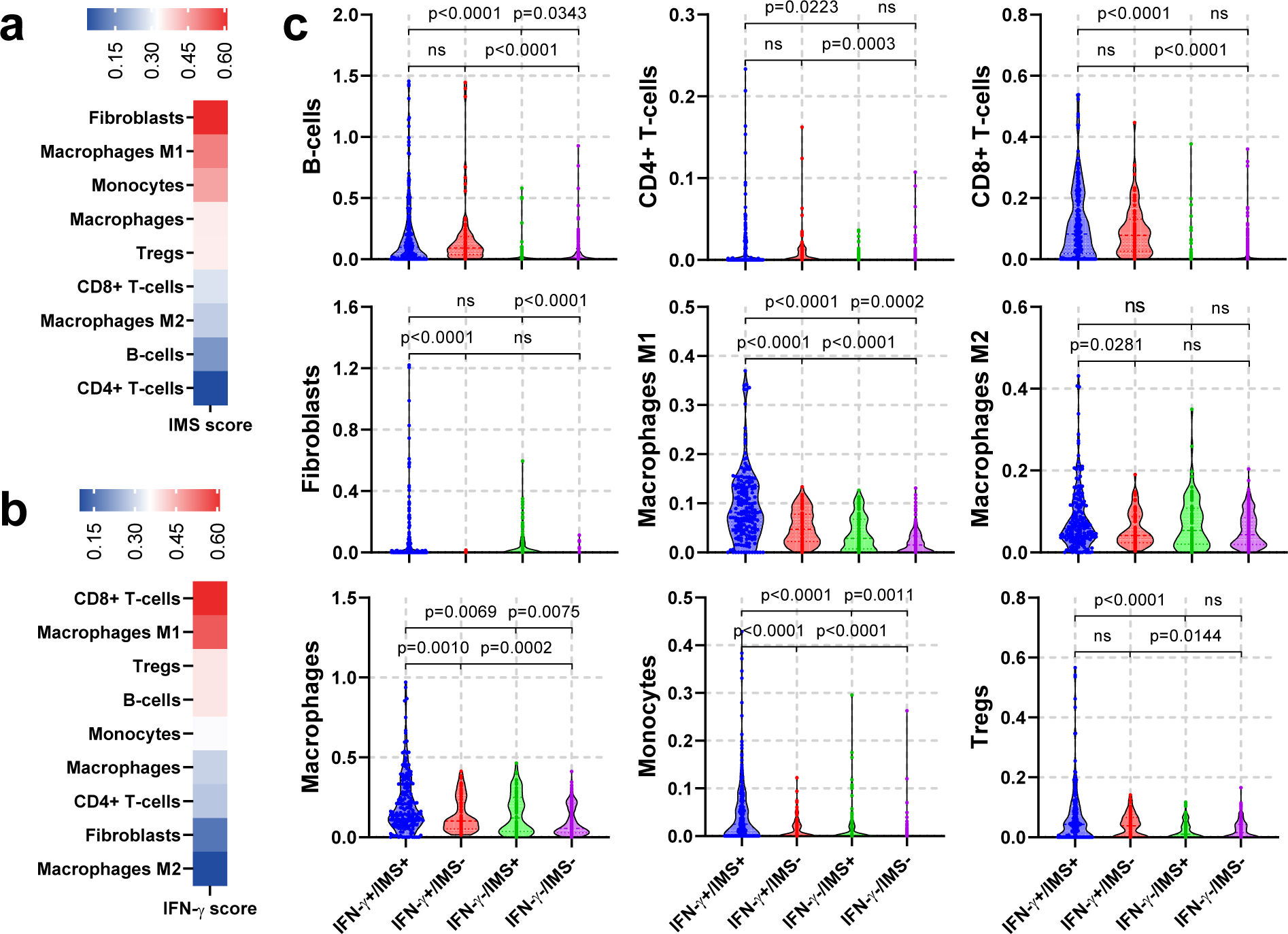
Association of IMS with abundance of immune cell types in TME. **a**-**b**, Heatmap showing the Pearson correlation of selected immune cell types with IMS scores (in **a**) and IFN-*γ* (in **b**) in combined melanoma cohorts consisting of all samples data from the discovery cohort and TCGA melanoma dataset (*n* = 516). Fibroblasts showed the strongest association with the IMS score (Pearson correlation *r* = 0.62, *p* < 0.001), followed by M1 macrophage (*r* = 0.50, *p* < 0.001), monocytes (*r* = 0.45, *p* < 0.001) and macrophage (*r* = 0.35, *p* < 0.001). On the other hand, CD8+ T-cell showed the strongest association with the IFN-*γ* score (*r* = 0.61, *p* < 0.0001). **c**, Violin plot showing the cell type distributions estimated using xCell for 4 groups of patients with IFN-*γ*+/IMS+ (*n* = 183), IFN-*γ*+/IMS- (*n* = 75), IFN-*γ*-/IMS+ (*n* = 75), IFG- *γ*-/IMS- (*n* = 183) in the combined melanoma cohort. Cutoff values for the IFN-*γ* signature and IMS scores were set to their median in all the data.

### Balance between the IFN-*γ* signature and the IMS as a biomarker for cancer

We next studied the distribution of IMS scores and their interaction with IFN-*γ* signature scores in different tumour types using TCGA data. First, we analysed the correlation between IMS scores and IFN-*γ* signature scores for all TCGA patients (*n* = 11, 043; Fig. 3a). The results showed that the IFN-*γ* signature and IMS scores had a modest positive correlation with *r* = 0.53 (*p* < 0.0001); however, IMS scores were poorly explained by IFN-*γ* signature scores (*R*^2^ = 0.28), suggesting that these two signatures are not fully overlapping and might contribute complementary information regarding the TME. A similar conclusion can be made on the correlation of the IMS and IFN-*γ* scores on selected cancer types from TCGA (Supplementary Fig. 1).

**Fig. 3.**
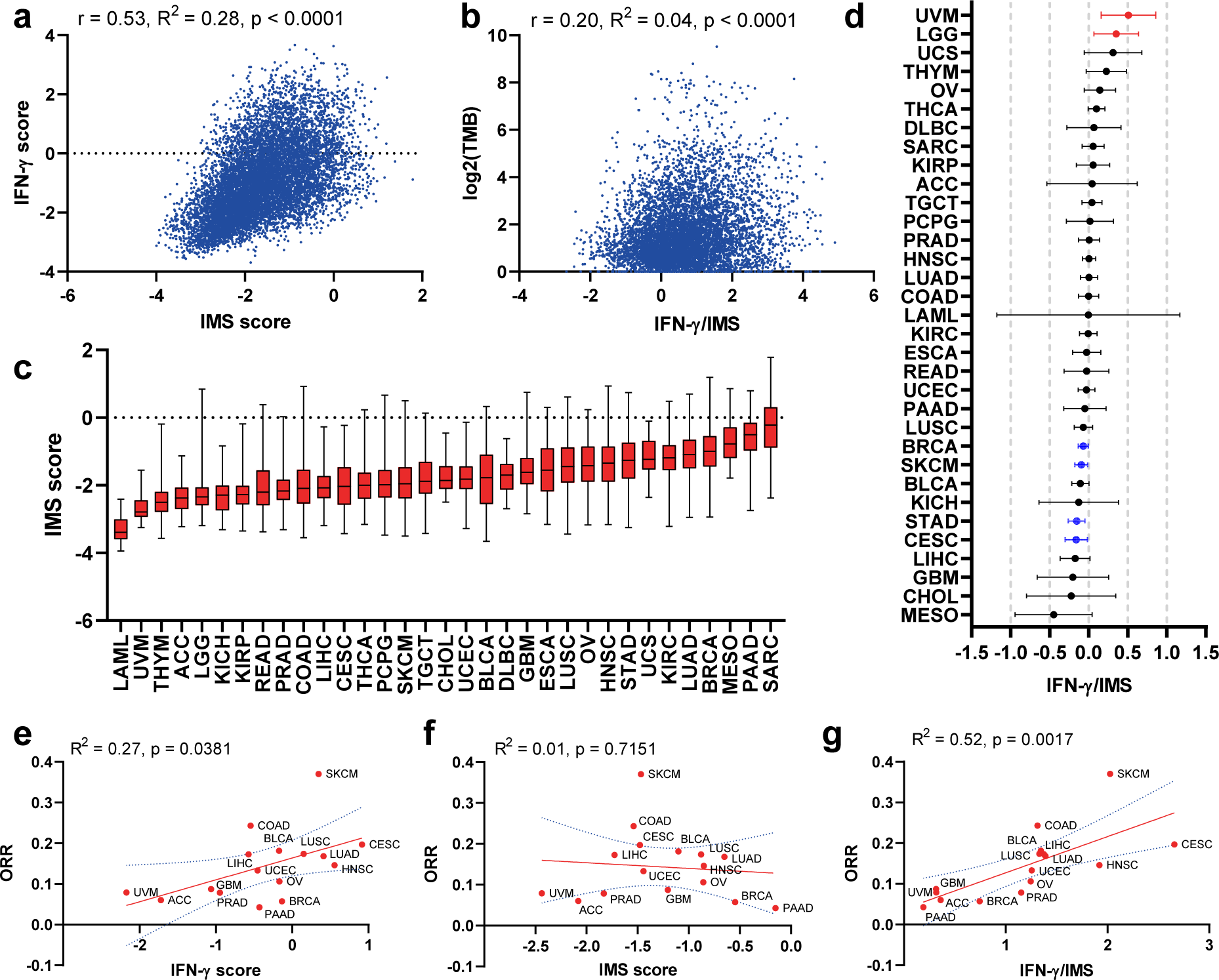
Balance between the INF-*γ* signature and IMS as a biomarker for cancer. **a**, Pearson correlations of IMS score with the IFN-*γ* signature (*r* = 0.54, *R*^2^ = 0.28, *p* < 0.0001) for all TCGA patients (*n* = 11, 043). **b**, Pearson correlation of logTMB with IFN-*γ*/IMS ratio (*r* = 0.20, *R*^2^ = 0.04, *p* < 0.0001) for all TCGA patients (*n* = 11, 043). **c**, Boxplots showing a summary of the distribution of IMS scores for all TCGA patients, with tumour types ordered by their median IMS score. **d**, Log hazard ratio estimates and 95% confidence intervals, with adjustment for sex, age, and TMB, with a binary cutoff (top 20% of each cancer type). Cancers in which the IFN-*γ*/IMS ratio was statistically significantly (*p* < 0.05) associated with good prognosis are highlighted in blue; significant associations with poor prognosis are in red. **e**-**g**, Associations of the ORR to immunotherapy for different cancer types with their median IFN-*γ* signature (linear regression goodness-of-fit *R*^2^ = 0.27, *p* = 0.381; in **e**), IMS (*R*^2^ = 0.01, *p* = 0.715; in **f**), and IFN-*γ*/IMS (*R*^2^ = 0.52, *p* = 0.017; in **g**) values in the TCGA datasets.

Given that the status of the tumour immune microenvironment and the associated composition of immune cells contain prognostic information, we hypothesized that the balance between IFN-*γ* signature and IMS may be associated with the survival of cancers. To assess this possibility, we performed a stratified multivariate analysis using Cox proportional hazards regression within each TCGA cancer type. The results showed that the association between the ratio of IFN-*γ* signature to the IMS score (IFN-*γ*/IMS) and overall survival (OS) varied according to cancer type (Fig. 3b). A higher IFN-*γ*/IMS ratio was associated with a modest prognostic benefit after adjusting for sex, age, and TMB in breast invasive carcinoma (BRCA) (HR = 0.93; 95% CI: 0.88 - 0.99), cutaneous melanoma (SKCM) (HR = 0.91, 95% CI: 0.84 - 0.98), stomach adenocarcinoma (STAD) (HR = 0.85; 95% CI: 0.76 - 0.94), and cervical tumours (CESC) (HR = 0.83; 95% CI: 0.73 - 0.96). Conversely, a higher IFN-*γ*/IMS ratio was associated with poor prognosis in uveal melanoma (UVM) (HR = 1.50; 95% CI: 1.07 - 2.11) and brain lower-grade gliomas (LGG) (HR = 1.41; 95% CI: 1.06 - 1.88), suggesting that these cancers may have different antitumour immune responses than those cancers mentioned previously. Interestingly, it was previously reported that a higher TMB was associated with poor survival in patients with glioma^7^. Associations of the IFN-*γ*/IMS ratio and survival in different directions have also been observed in other cancer types but did not reach statistical significance.

We further checked the relationship between the IFN-*γ*/IMS ratio and TMB scores in TCGA data and found that there was a positive but weak association between them (*r* = 0.20, *R*^2^ = 0.042, *p* < 0.0001; Fig. 3b) on all samples from TCGA datasets (*n* = 11, 043) and on selected cancer types (Supplementary Fig. 1).

Finally, we compared the median values of the IFN-*γ* signature score, the IMS score, and their ratio with objective response rates (ORRs) to anti-PD-1 therapies for cancer types with efficacy performance data available^31^. A positive correlation of ORR with the IFN-*γ* signature score

(*R*^2^ = 0.27, *p* = 0.047, Fig. 3e) and IFN-*γ*/IMS (*R*^2^ = 0.54, *p* = 0.001, Fig. 3g) was observed (Fig. 3f). Importantly, tumours with high median IFN-*γ*/IMS values, most notably SKCM^32^, colon adenocarcinoma (COAD), CESC^33^, bladder urothelial carcinoma (BLCA)^34^, lung squamous cell carcinoma (LUSC)^35^ and liver hepatocellular carcinoma (LIHC), have shown clinical sensitivity to ICI therapies (Fig. 3g). Some tumour types (e.g., pancreatic adenocarcinoma (PAAD) and BRCA) have shown poor responses to immunotherapy despite their moderate to high median IFN-*γ* scores (Fig. 3e). These tumours are known to be highly infiltrated with myeloid cells, which may serve as an additional immunosuppressive mechanism preventing efficacy with ICI therapy^36, 37^. Notably, these cancer types were also characterized by elevated IMS scores (Fig. 3c), and hence, a better association with ORR to ICI therapies was observed when both signatures were considered.

### Ratio of the IFN-*γ* signature score to the IMS score predicts PD-1 blockade efficacy

We next assessed whether directly using the ratio of the IFN-*γ* signature score to the IMS score could be used as a reliable metric to predict anti-PD-1 therapy outcome for melanoma patients. We used IFN-*γ*/IMS together with the clinical response data to generate receiver operating characteristic (ROC) curves to quantify its prediction performance in our discovery cohorts. The resulting AUCs were in the range of 0.70 - 0.83 (Supplementary Fig. 3b).

We next tested the prediction ability of IFN-*γ*/IMS in a newly generated RNA-seq dataset from 55 tumour tissues of melanoma patients treated with anti-PD-1 monotherapy at Peking University Cancer Hospital (PUCH), Beijing, China. In this dataset, IFN-*γ*/IMS achieved a prediction accuracy of AUC = 0.81 (95% CI: 0.69 - 0.93; Fig. 4e). Using the threshold that generated the maximum Youden index^38^ to divide patients into predicted responder (*n* = 29) and predicted non-responder groups (*n* = 26), IFN-*γ*/IMS successfully captured 67.71% of nonresponders (23 out of 35) with only 3 exceptions (1 patient with a PR/CR and 2 patients with SD were misclassified as nonresponders), achieving a classification accuracy of 88.5% (*p* = 0.0006) for this group. On the other hand, of the 29 patients classified as predicted responders by IFN-*γ*/IMS, 17 (13 patients with a PR/CR and 4 patients with SD) actually responded. Overall, a higher IFN-*γ*/IMS ratio was associated with a better ORR (*p* = 0.0005; Fig. 4a), OS (HR = 2.78; 95% CI: 0.80 - 9.64; *p* = 0.1214; Fig. 4h) and progression free survival (PFS) (HR = 3.45; 95% CI: 0.97 - 12.19; *p* = 0.0547; Fig. 4g) than a lower IFN-*γ*/IMS ratio. Compared with IFN-*γ* signature-based classification (Fig. 4b), IFN-*γ*/IMS correctly classified six nonresponders that would otherwise be misclassified by IFN-*γ* signature as responders due to their medium to high IFN-*γ* scores, and one patient who had low IFN-*γ* score but responded to ICI therapy as responder (Fig. 4c). The OS and PFS results were not significant due to limited sample size, and relatively short follow-up period of this cohort. However, it was recently found that OS and PFS were significantly different between clinical responders and progressors of melanoma patients to anti-PD-1 therapy^13^, suggesting that short-term response could be used as a surrogate for the survival benefits of patients in this context. In the public dataset of 48 preclinical metastasis melanoma treated with anti-PD-1 (Liu19^13^), IFN-*γ*/IMS achieved an AUC of 0.66 (95% CI: 0.50 - 0.83, Fig. 5b). In addition, patients with higher IFN-*γ*/IMS ratio (with cutoff value based on the Youden index) had better ORR (*p* = 0.0043; Supplementary Fig. 2e) and longer PFS (HR = 4.36; 95% CI: 1.36 - 14.03; *p* = 0.0013, Fig. 5d). Collectively, the above data demonstrate the potential value of IFN-*γ*/IMS ratio as a combinatorial biomarker for anti-PD-1 treatment for metastastic melanoma.

**Fig. 4.**
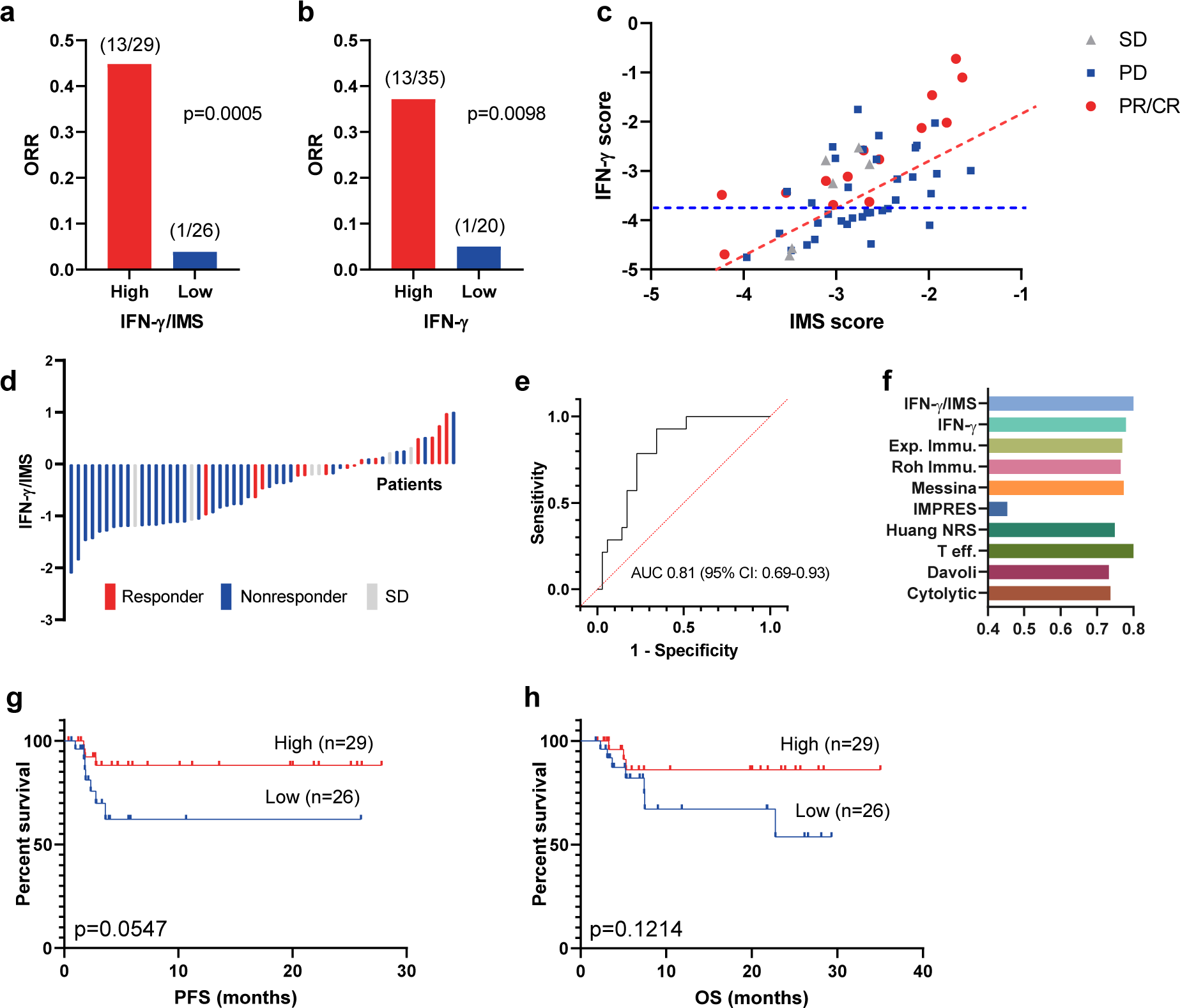
Ratio of IFN-*γ* signature and IMS predicts response to ICI immunotherapy on the PUCH cohort. **a**, The ORR of patients from IFN-*γ*/IMS-high vs IFN-*γ*/IMS-low and **b** IFN-*γ*-high vs IFN-*γ*-low on the PUCH cohort. The cutoff points were decided by the Youden index for IFN-*γ*/IMS and IFN-*γ* scores, respectively. **c**, IFN-*γ* signature and IMS scores of individual patients in the PUCH cohort. The red and blue dashed lines indicate the cutoff points for IFN-*γ*/IMS ratio and IFN-*γ*, respectively. **d**, Waterfall plots of IFN-*γ*/IMS versus patients with different clinical responses to anti-PD-1 therapy in the PUCH cohort. **e**, ROC curve of the sensitivity versus 1-specificity of the predictive performance of IFN-*γ*/IMS. Patients with SD were not included in AUC calculation. **f**, Comparison of the AUC of IFN-*γ*/IMS with nine GEP signatures (Table 2) in predicting response to ICI. **g**, Kaplan-Meier plots of OS and PFS segregated by IFN-*γ*/IMS ratio with cutoff points selected according to the Youden index.

**Fig. 5.**
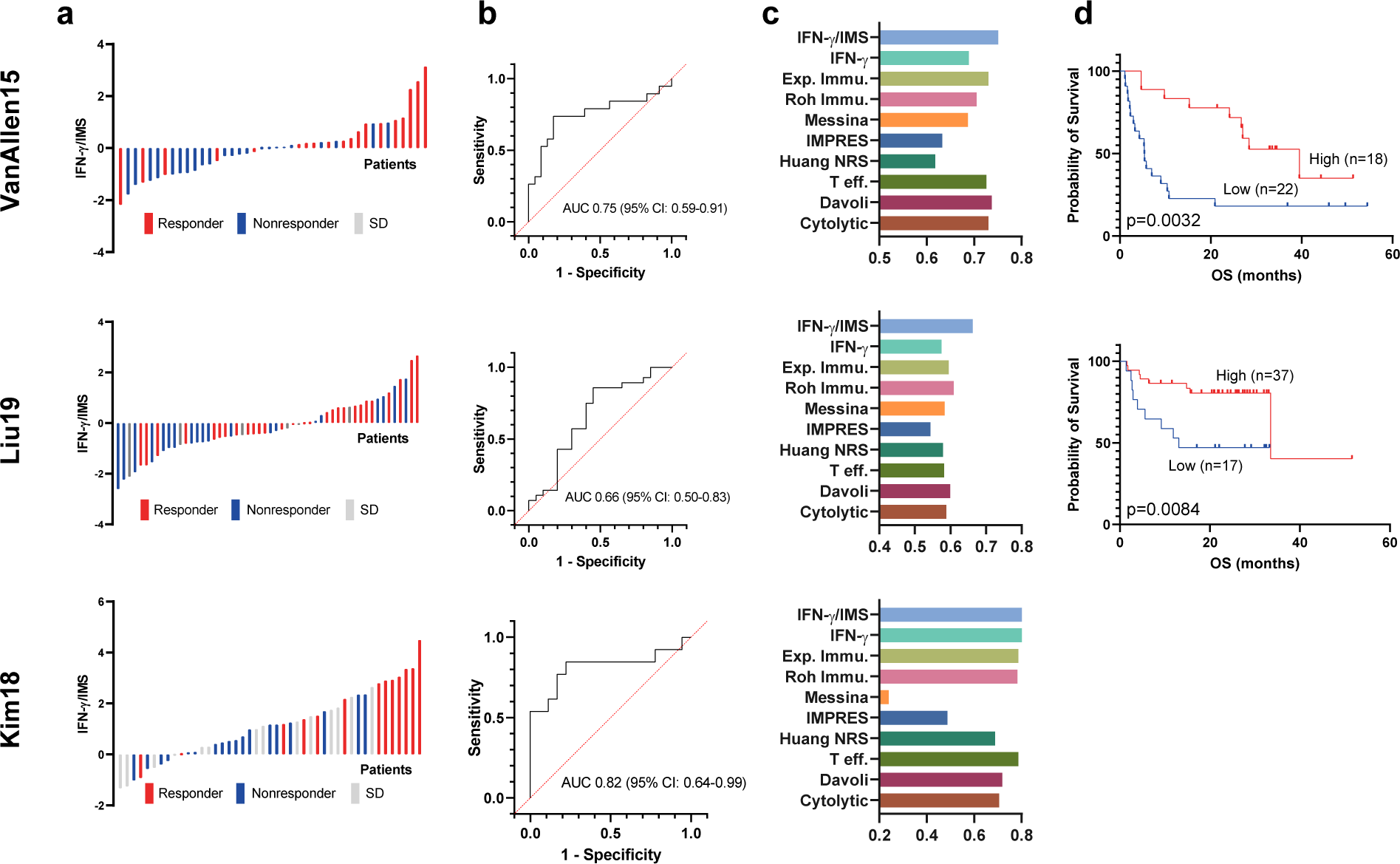
Ratio of IFN-*γ* signature and IMS predicts response to ICI immunotherapy on published datasets. **a**, Waterfall plots of the IFN-*γ*/IMS versus patients with different clinical responses to anti-PD-1 therapy in each validation cohort. **b**, ROC curve of the sensitivity versus 1-specificity of the predictive performance of IFN-*γ*/IMS. Patients with SD were not included in AUC calculation. **c**, Comparison of the AUC of IFN-*γ*/IMS with nine GEP signatures (Table 2) in predicting response to ICI. **d**, Kaplan-Meier plots of OS or PFS segregated by IFN-*γ*/IMS ratio with cutoff values selected according to the Youden index identified in individual datasets. Kaplan-Meier plot for Kim19 was not generated due to unavailability of survival data for this dataset.

Although the IMS was derived from advanced melanoma cohorts receiving anti-PD-1 treatment, the resulting genes measure immune-related expression levels with minimal contribution from tumour-related transcriptomic activities. Therefore, the signature may provide a treatment or tumour-type agnostic insight into immune microenvironment activities. To test this concept, we further evaluated the prediction performance of IFN-*γ*/IMS on two publicly available RNA-seq datasets with pretreatment samples from melanoma patients treated with anti-CTLA4 therapy (VanAllen15; *n* = 42)^39^ and metastatic gastric cancer patients treated with anti-PD-1 therapy (Kim18; *n* = 45)^40^. The resulting AUCs were 0.75 (95% CI: 0.59 - 0.91; Fig. 5b) and 0.82 (95% CI: 0.64 - 0.99; Fig. 5b), respectively, for these two datasets. In addition, patients with high IFN-*γ*/IMS ratios (using the Youden index to determine the cutoff point) had better ORR on the VanAllen15 dataset (*p* = 0.0004; Supplementary Fig. 2d) and the Kim18 dataset (*p* = 0.0022; Supplementary Fig. 2f) and longer OS in the VanAllen15 dataset (HR = 3.06; 95% CI: 1.41 - 6.61; *p* = 0.0032; Fig. 5d) than patients with low IFN-*γ*/IMS ratios, suggesting the potential of using the IFN-*γ*/IMS ratio as a predictive/prognostic biomarker for immunotherapies different to anti-PD-1, or other cancer types.

**Table 1:** Cohorts used in this study

| Cohort name | Tumour type | Cohort size | Target checkpoint |
| --- | --- | --- | --- |
| PUCH | Melanoma | 55 | PD-1 |
| Riaz17<sup>15</sup> | Melanoma | 51 | PD-1 |
| Gide19<sup>21</sup> | Melanoma | 41 | PD-1 |
| Hugo16<sup>14</sup> | Melanoma | 28 | PD-1 |
| VanAllen15<sup>39</sup> | Melanoma | 42 | CTLA-4 |
| Liu19<sup>13</sup> | Melanoma | 54 | PD-1 |
| Kim18<sup>40</sup> | Gastric | 45 | PD-1 |

**Table 2:** GEP signatures used in this study

| Signature name | Number of genes | Description |
| --- | --- | --- |
| IFN-γ/IMS | 28 | This work. |
| IFN-γ<sup>43</sup> | 6 | Averaging the expression levels of the IFN-γ signature genes. |
| Exp. Immu.<sup>43</sup> | 18 | Averaging the expression levels of the expanded immune genes. |
| Roh Immu.<sup>77</sup> | 41 | Averaging the expression levels of immune genes. |
| Messina<sup>78</sup> | 12 | Principal component 1 score from PCA of expression levels of 12 chemokine signature genes. |
| IMPRES<sup>79</sup> | 28 | Sum of ratios of 15 checkpoint or immune gene pairs. |
| Huang NRS<sup>80</sup> | 69 | Averaging the expression levels of neoadjuvant response signature (NRS) genes. |
| T eff.<sup>81</sup> | 8 | Averaging the expression levels of T-effector IFN-γ signature genes. |
| Davoli<sup>82</sup> | 7 | Averaging the expression levels of cytotoxic immune signature genes |
| Cytotoxic<sup>83</sup> | 2 | Averaging the expression levels of granzyme A (GZMA) and perforin (PRF1). |

### Comparison with other GEP signatures

Currently, there were significant number of independent studies on GEP signatures that predict the response of patients to anti-PD-1 therapy. To compare the prediction accuracy of the proposed IFN-*γ*/IMS ratio with that of existing GEP signature-based predictors, we generated predictions using nine published GEP signatures (Table 2) and the comparison results showed that IFN-*γ*/IMS ratio achieved better AUC performance on both PUCH (Fig. 4f) and the three external validation cohorts (Fig. 5c). One limitation of existing GEP signature studies is that many of these signatures were validated with independent cohorts within each publication, but frequently these signatures have not performed well in follow-up reports. To further validate the robustness of the proposed approach, we performed a randomized permutation test where three datasets were randomly selected from the seven datasets (Table 1) as the discovery cohort to identify the top 18 IMS genes as described previously. We then tested the prediction performance of ratio of IFN-*γ* and the identified IMS on the remaining four datasets. The results from the total 35 purmutation tests indicated that IFN-*γ*/IMS outperformed other GEP singatures by a significant margin (Wilcoxon matched-pairs signed rank test, *p* < 0.0001; Fig. 6a). Significantly, we found that the IMS signatures from the randomized tests were highly consistent despite that they were obtained from different training datasets. More than half of the total 630 occurrences of the IMS genes from the 35 randomized tests were from the top 23 frequent genes (Fig. 6b). Moreover, of the 18 IMS genes identified from the original discovery cohorts, 13 (OLFML2B, AXL, ADAM12, STC1, VCAN, PDGFRB, INHBA, CAT1, COL6A3, SIGLEC1, CD163, IL10, TWIST2) can be found in these top 23 frequent IMS genes from the randomized test. Further analysis of these genes on a public single-cell RNA-seq (scRNA) dataset from melanoma^41^ indicated that most of these genes are highly expressed on CAF (e.g., OLFML2B, VCAN, PDGFRB, COL6A3; Fig. 6d and Supplementary Fig. 4) and/or macrophages (e.g., VCAN, CD163, SIGLEC1; Fig. 6d and Supplementary Fig. 4), confirming the significant roles of these immune cells and their related immune suppressive activities in preventing patients from responding to anti-PD-1 therapy.

**Fig. 6.**
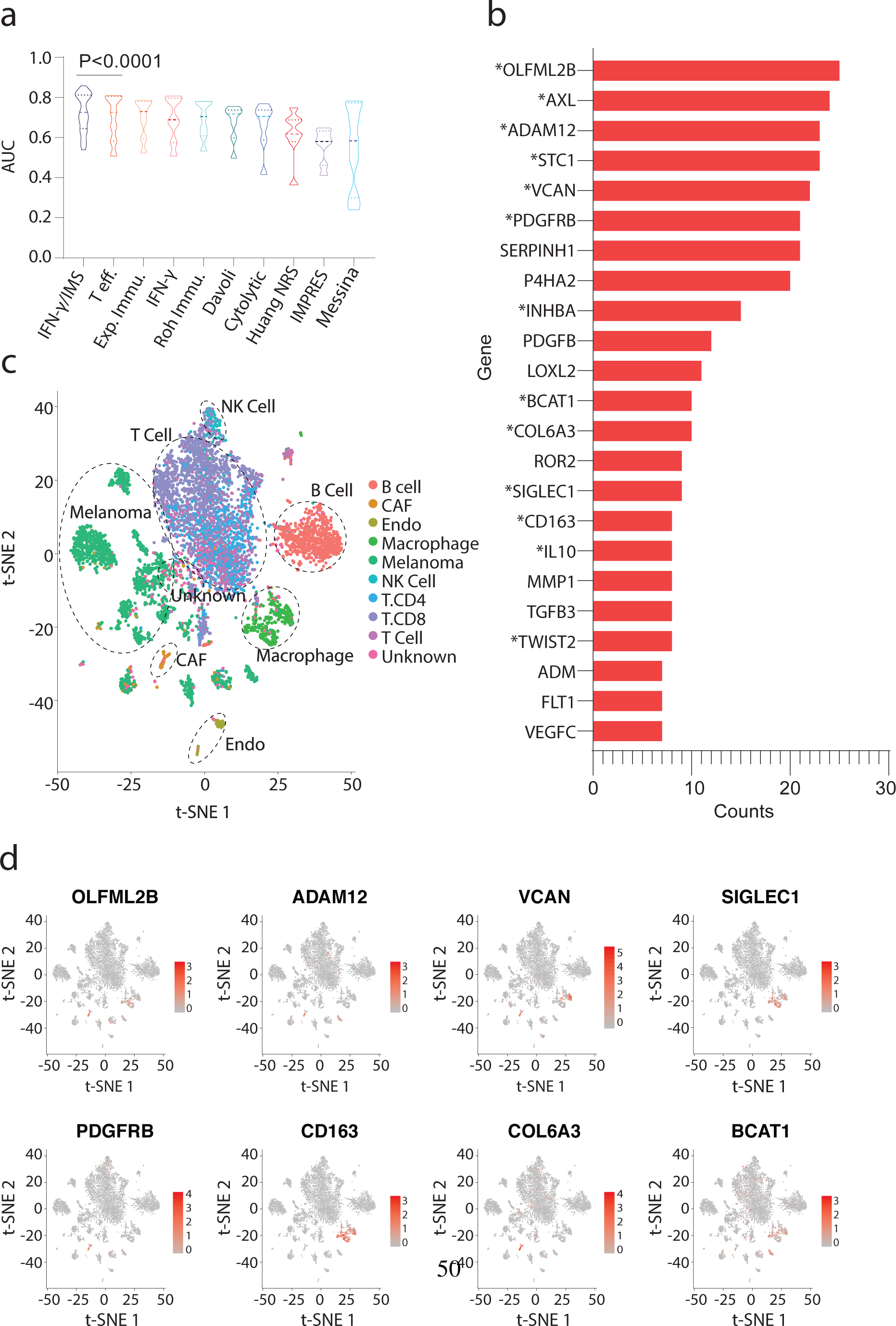
Robustness of the proposed IMS signatures. **a**, Comparison of AUC performance of IFN-*γ*/IMS ratio with nine published GEP signatures (Table 2) after 35 bootstrapping randomized tests on the seven datasets (Table 1). In each randomized test, the IMS genes were identified from three randomly selected datasets (Methods), and the AUC prediction performance of the ratio of IFN-*γ* signature and the IMS identified from the randomly selected dataset was evaluated on the remaining four datasets. Models are sorted by their median AUC performances and Wilcoxon matched-pairs signed rank test was performed to compare the AUC performances of IFN-*γ*/IMS and the second best model T eff. **b**, Top 23 highly frequent genes from the bootstrapping tests. Genes in the original IMS signature are marked with an asterisk (*) symbol. **c-d**, t-SNE plot of cells from melanoma^41^. Cells are colored by cell types in (**c**) and by normalized expression of different IMS genes in (**d**).

## 3 Discussion

There is significant interest in developing robust biomarkers of response to immunotherapy, as well as identifying actionable targets in those who do not respond to the current standard ICI therapies. Gene expression biomarkers, such as Oncotype DX^42^, have demonstrated clinical utility in predicting treatment benefits in breast cancer. However, as interactions between the tumour and its microenvironment are highly complex, constructing predictors of patient response to ICIs remains a serious challenge.

Existing efforts to create gene expression-based tests for ICI efficacy have mainly focused on developing “response signatures” that measure the expression of adaptive immune response-related inflammatory genes^20^, most of which include an IFN-*γ* gene signature as a major component^43^. However, due to the presence of intricate immunosuppressive mechanisms within the tumour microenvironment (TME), the presence of a peripherally suppressed adaptive immune response alone appears to be necessary but not sufficient for clinical benefit from PD-1/PD-L1 blockade. In this study, we identified an immunosuppression signature that, when combined with an inflammatory signature, had prognostic/predictive value in patients with advanced melanoma treated with PD-1 blockade. To maximize our chance in identifying the correct genes for this signature, we started with an established 10-gene IFN-*γ* signature measuring the expression of genes associated with cytotoxic cells, antigen presentation, and IFN-*γ* activity^43^ and then selected genes that were consistently upregulated in nonresponsive vs responsive groups after adjusting for the IFN-*γ* signature score based on their *p*-values on multiple datasets. In addition, since conceptually, a truly predictive gene should produce a significant result in all datasets, we used Pearson’s method^44^, which is more sensitive to the largest *p*-value, to combine *p*-values from different datasets to avoid artefacts due to single significance from individual dataset^45^.

Strikingly, the genes identified in our IMS through the above computational method were highly consistent with several important biological activities related to innate or acquired resistance to ICIs. CAFs are a nonredundant, immunosuppressive component of the TME^46, 47^. It was previously reported that INHBA production by cancer cells helps to induce CAFs, and ablating inhibin *β* A decreases the CAF phenotype both in vitro and in vivo^48^. CAFs hinder antitumour immunity by secreting immunosuppressive cytokines such as IL-10 and TGF-*β*, reducing CTL function and viability^49, 50^ and attracting immunosuppressive myeloid cells, including tumour-associated macrophage (TAMs), via CCL2^47, 51^. Notably, SIGLEC1/CD163 is associated with the activation of macrophages towards an immunosuppressive phenotype, and accordingly, the expression of both CAF (FAP+) and TAM (SIGLEC1+) markers is associated with poor clinical outcomes across multiple tumour types^52–55^. CAF-mediated EMT, which is strongly correlated with the expression of AXL, TWIST2 and ADAM12 from the IMS, can result in biomechanical and biochemical changes that facilitate tumour immune escape, invasion, and metastasis^56^. The dense collagen matrix produced by CAFs may also present a physical barrier to the infiltration of T lymphocytes^57^ or treatments reaching the cancer cells^58^. Indeed, the association between a lack of response to ICIs and upregulated EMT-related genes has been observed in multiple cancers^40, 59^, and inhibiting CAF/TAM-related pathways and extracellular collagen and hyaluronan can induce T cell accumulation and improve the outcome of ICIs^60–63^, reinforcing the role of those stromal-related activities in limiting the efficacy of immune checkpoint blockade immunotherapy.

Using the ratio of opposing immune signatures instead of the absolute value of individual signatures as a predictive/prognostic biomarker brings another advantage. It is well known that to compensate for potential technical variation, raw gene expression data from RNA-seq must be normalized so that meaningful biological comparisons can be made^64^. Typically, this is done with a set of housekeeping genes that are expected to maintain constant expression levels under different experimental conditions^65^. However, it has become increasingly clear that housekeeping gene expression levels may vary considerably in some conditions^66, 67^. When that happens, the normalization process itself can lead to increased intersample “noise” that covers meaningful differences in target genes if the chosen housekeeping genes fluctuate randomly or erroneous results if there is a directional change in the housekeeping genes between experimental groups^66, 68^. The calculation of the IFN-*γ*/IMS ratio provides a self-normalization method that directly measures the balance between contradicting biological processes within the tumour microenvironment, thus eliminating the need for using housekeeping genes that can be unreliable.

Our study has limitations. Since the current ICI clinical trials have generated gene expression data for only a limited number of pretreatment samples, which were insufficient to train robust prediction models, we did not systematically optimize the weights of individual genes in the IFN-*γ*/IMS ratio calculation. With more RNA-seq data available from subsequent studies, we expect that further optimization of the combined biomarker will yield even better predictive accuracy. In this study, we did not attempt to specify a universally applicable cutoff point for IFN-*γ*/IMS for different datasets due to potential batch effects from different RNA-seq procedures conducted at multiple sites. Rather, we demonstrated a trend that shows an increase in benefits with increasing IFN-*γ*/IMS ratios. Nevertheless, we envision that a relevant cutoff would need to be aligned to specific assay designs and clinical situations, and such a cutoff point could be further standardized based on additional evidence of merits from future clinical studies. Finally, our analysis is retrospective in nature, and validation of the findings in additional datasets is warranted.

In conclusion, the IMS studied in this paper exemplifies the potential of using GEP signatures for modeling the adverse TME, and using IMS in combination with an existing inflammatory GEP signature enables better identification of patients who could respond favorably to ICIs. Currently, clinical trials are assessing the efficacy of combining anti-PD-1 therapy with medicines that target at normalization of immune suppressive TME including the CSF1R inhibitor Cabrilizumab for the treatment of resectable biliary tract cancer (NCT03768531), CCR2 inhibitor plozalizumab for the treatment of melanoma (NCT02723006), the FAP inhibitor RO6874281 for the treatment of metastatic head and neck, oesophageal or cervical cancers (NCT03386721) and metastatic melanoma (NCT03875079), and the TGF-*β* inhibitor galunisertib (LY2157299) for the treatment of advanced-stage NSCLC or hepatocellular carcinoma (NCT02423343) and metastatic pancreatic cancer (NCT02734160). However, due to the diversity of the immune evasion mechanisms in inflammatory tumours, such as loss of heterozygosity at the HLA locus^69^, mutations in JAK-STAT signalling^70^ or loss of IFN-*γ* pathway genes^71^, reduced expression or loss of function of *β*2-microglobulin^72^, and loss of immunogenic mutations^73^ or epigenetic repression of neoantigen transcripts^74^, specific immunosuppressive mechanisms utilized by each individual tumour would still need to be fully understood and gauged to better direct patients to different combination therapy options. In this regard, it is anticipated that IMS or future immunosuppressive signatures gleaning through deeper understanding of the immunosuppressive mechanisms of cancer would enable the development of more effective stratification models or therapeutic combinations to increase the efficacy and cost-effectiveness of immunotherapies for the benefits of cancer patients.

## 4 Methods

### Patients and tissue samples

In this study, we obtained 55 formalin-fixed, paraffin-embedded (FFPE) tumour tissues from melanoma patients treated with anti-PD-1 monotherapy at PUCH, Beijing, China, between March 2016 and March 2019 (Supplementary Table 2). Diagnosis was histopathologically confirmed for all patients. Clinical data, including sex, age, tumour site, tumour thickness, metastasis status, and clinical efficacy, were collected. Therapy outcomes evaluated following Response Evaluation Criteria in Solid Tumors (RECIST) version 1.1, including presence of a complete or partial response (CR/PR), stable disease (SD) and progressive disease (PD), were used to assess efficacy. OS was calculated from the treatment start date. Patients who did not die were censored at the date of last contact.

### Whole-transcriptome RNA sequencing

Total RNA was extracted from unstained FFPE tumour samples by the All Prep-DNA/RNA-Micro Kit (Qiagen) following the standard manufacturer’s protocol. Reverse transcription and second-strand cDNA synthesis were subsequently performed. Barcoded RNA libraries were generated and captured by a customized whole-exome panel. All libraries were sequenced on the Illumina NovaSeq 6000 platform with 2×150 bp paired-end reads. The mean sequencing coverage across all samples was ∼100X (3.5 G). RNA-seq reads were mapped to the human reference genome GRCh37 using STAR^75^, and gene expression was quantified using RSEM^76^. Coding region reads were counted to calculate fragments per kilobase of transcript per million mapped reads (FPKM) values at the gene level and log2-transformed before analysis to avoid extremely skewed gene expression distributions.

### External data sources

We collected the RNA-seq data of melanoma patients from six immunotherapy studies with gene expression profiles for pretreatment tumours and complete clinical information, including the Riaz17 (*n* = 51)^15^, Hugo16 (*n* = 28)^14^, Gide19 (*n* = 41)^21^, VanAllen15 (*n* = 42)^39^, Liu19 (*n* = 54)^13^ and Kim18 (*n* = 45)^40^ datasets (Table 1). Patients from these clinical studies were treated with nivolumab^15^ and/or pembrolizumab^14, 21^. For the Gide19 and Liu19 studies, only baseline data from samples that received anti-PD-1 monotherapy (nivolumab or pembrolizumab) were used. The immunotherapy outcomes provided in the original publications following RECIST guidelines (PR/CR/SD/PD) were used in our analysis. The gene expression data of VanAllen15 and Liu19 are download from respective references as provided by the authors. For Riaz17, Hugo16, Gide19 and Kim18, the RNA-seq raw data was obtained and processed by the above mentioned pipeline to generate the gene expression data.

We downloaded TCGA Level-3 RSEM-normalized RNA-seq data and mutation packer calls from the TCGA database. The RNA-seq data were log2-transformed. Each patient’s TMB was calculated as the number of nonsynonymous mutations.

### Housekeeping normalization

We renormalized the RNA-seq data using a set of 20 reference (“housekeeping”) genes (ABCF1, DNAJC14, ERCC3, G6PD, GUSB, MRPL19, NRDE2, OAZ1, POLR2A, PSMC4, PUM1, SDHA, SF3A1, STK11IP, TBC1D10B, TBP, TFRC, TLK2, TMUB2, and UBB) with low variance across a set of banked tumour samples from a variety of cancer types. The log2-transformed expression of each gene was normalized by subtracting the arithmetic mean of the log2-transformed expressions of the housekeeping genes.

### Identification of the IMS genes

To identify the IMS genes, we performed a one-sided Student’s *t*-test to capture genes that were systematically upregulated in the nonresponse groups (PD) vs response groups (PR/CR) after normalization by the IFN-*γ* signature score in each individual dataset. Due to the large dimensionality of the data, we restricted our search to the 770 cancer immune-related genes curated in Nanostring’s IO 360 panel. The resulting *p*-values from the three datasets in the discovery cohort were combined using Pearson’s method^44^ to avoid artefacts due to single significance from individual dataset. The genes were ranked based on their Pearson combined *p*-values, and the top 18 genes were identified as our IMS genes.

### Calculation of GEP signatures

We collected nine published GEP signatures related to the immune checkpoint response from the literature and validated in our cohorts (Table 2). Sample-wise scores of these signatures were calculated from RNA-seq data following the methodology described in the corresponding papers. Genes with unavailable expression data were excluded from the calculation of signature scores.

For the IFN-*γ* signature and IMS scores in this paper, we used the arithmetic mean of the log2-transformed, housekeeping gene normalized expression level of the 10-gene “preliminary” IFN-*γ* signature (IFNG, STAT1, CCR5, CXCL9, CXCL10, CXCL11, IDO1, PRF1, GZMA, and HLA-DRA)^43^, and the 18 IMS genes listed in Supplementary Table 1 respectively. Furthermore, the IFN-*γ* signature/IMS ratio was calculated as the difference between these two scores in the logarithmic domain.

### Single cell RNA-seq

Briefly, scRNA-seq data of 31 melanoma tumors were downloaded from GEO database (GSE115978)^41^. The original expression profiles and cell type annotations were used. Principal component analysis (PCA) was performed to reduce the dimensionality of the scRNA-seq profiles. Then t-SNE projections were generated using the first 25 principal components. Both PCA and t-SNE analysis were performed by RunPCA and RunTSNE functions in the Seurat package (version 3.1.0) with default parameters.

### Data analysis and statistical information

Associations between categorical measurements and patient groups, such as the predictive accuracy of different biomarkers/panels, were evaluated using Fisher’s exact test. Differences in continuous measurements were tested using the two-tailed Mann-Whitney U-test. Correlations between two groups of continuous variables were evaluated using Pearson correlation analysis. The Kaplan-Meier method was utilized to estimate overall survival, and difference between groups were assessed using the log-rank test. Two-sided *p*-values were used unless otherwise specified, and a *p*-value less than 0.05 was considered significant. For boxplots, centre mark is median and whiskers are minimum/maximum unless specified otherwise.

PRISM was used for basic statistical analysis and plotting (http://www.graphpad.com), and the Python language and programming environment were used for the remainder of the statistical analysis. The abundances of multiple cell types in whole tissue samples were estimated using xCell^30^.

### Code availability

Codes are implemented in Python and are publicly available in GitHub: http://github.com/xxx.

### Data availability

All patients data analysed from published papers are referenced to and publicly available accordingly. The gene expression data of patients from PUCH cohort will be deposited in the NCBI database once the manuscript is accepted for publishing. All the other data supporting the findings of this study are available within the article and its Supplementary Information files and from the corresponding author upon reasonable request.

## DECLARATIONS

### Study approval

The original studies were conducted in accordance with the Declaration of Helsinki and the International Conference on Harmonization Good Clinical Practice guidelines and approved by relevant regulatory and independent ethics committees from each study’s institution. All patients provided written informed consent before study entry.

### Authors’ contributions

RY and JG designed the study; YK, CX and WY performed the data processing and machine learning analysis. CC, XS, LS, YX and JY collected tumour samples of patients and pooled clinical and survival data from Peking University Cancer Hospital. XC and SW performed the experiments; WX, SY, JH and WZ contributed to the analysis of the data; All authors contributed to the drafting and the revision of the manuscript.

## Acknowledgements

We would like to thank Xu Xiao for her help in the analysis and visualization of the scRNA-seq data.

