## Supplementary for "Ratio of the interferon-*γ* signature to the immunosuppression signature predicts anti-PD-1 therapy response in melanoma"

### Supplementary material for "Ratio of the interferon-$\gamma$ signature to the immunosuppression signature predicts anti-PD-1 therapy response in melanoma"

Y Kong et al.

**Supplementary Fig. 1 | Correlation analysis between IMS and IFN-$\gamma$, IMS and TMB on selected TCGA datasets.** **a-h**, Pearson correlation analysis between IFN-$\gamma$ signature and IMS scores, and between IFN-$\gamma$/IMS ratio and TMB, on the BRCA, COAD, LUAD, and SKCM datasets from TCGA.

**Supplementary Fig. 2 | Comparison of ORR from IFN-$\gamma$/IMS-high and IFN-$\gamma$/IMS-low patients on individual cohorts.** **a-f**, Top: Comparison of ORR for patients from IFN-$\gamma$/IMS-high group versus patients from IFN-$\gamma$/IMS-low group with cutoff point (left), and for patients from IFN-$\gamma$-high group versus patients from IFN-$\gamma$-low group (right). Cutoff points were decided according to the Youden index on each cohort. Bottom: IFN-$\gamma$ signature and IMS scores of individual patients in each cohort. Red and blue dashed lines indicate cutoff points for IFN-$\gamma$/IMS and IFN-$\gamma$ signature score, respectively.

**Supplementary Fig. 3 | Ratio of IFN-$\gamma$ signature to IMS predicts response to ICI treatment on the discovery cohort.** **a**, Waterfall plots of the IFN-$\gamma$/IMS ratio versus patients with different clinical responses to anti-PD-1 therapy from different cohorts in the combined discovery cohort. **b**, ROC curve of sensitivity versus 1-specificity for the predictive performance of the IFN-$\gamma$/IMS ratio. The AUC values were 0.70 (95% CI: 0.50 - 0.90) for Hugo16, 0.83 (95% CI: 6.68 - 0.99) for

Gide19, and 0.76 (95% CI: 0.59 - 0.94) for Riaz17. Patients with SD were not included in AUC calculation. **c**, Comparison of the AUC of the IFN-$\gamma$/IMS ratio with nine GEP signatures in predicting response to ICI therapy from the literature: IFN-$\gamma$, Exp. Immu., Messina, IMPRES, T eff., Davoli, Cytolytic, Roh Immu., and Huang NRS. **d**, Kaplan-Meier plots of OS or PFS segregated by IFN-$\gamma$/IMS with cutoff points selected according to the Youden index on individual cohorts.

**Supplementary Fig. 4 | t-SNE plot of cells from melanoma.** Cells are colored by normalized expression of different IMS genes.

Supplementary Table. 1: Genes in the immunosuppression signature

| IMS biology | Gene | Putative function |
|---|---|---|
| Markers of immune cell | FAP, PDGFRB | CAFs |
| | CD163 | Tumor-associated macrophages |
| | SIGLEC1 (CD169) | Tissue resident macrophages |
| Cytokines | IL10 | Major immunosuppressive cytokine |
| | CCL2, CCL8, CCL13 | chemokines that recruit immunosuppressive cells |
| Stromal factors | INHBA | Contribute to immune escape of tumours by inducing CAFs |
| | VCAN | Recruit and activate immunosuppressive myeloid cells; associated with a decrease in tumour-infiltrating cytotoxic T cells |
| | AXL | Promote EMT, tumour angiogenesis and inhibit anti-tumor immune response |
| | TWIST2, ADAM12 | Promote tumour invasion and metastasis by inducing EMT and invadopodia-mediated ECM degradation |
| | COL6A3 | Increase proliferation and decrease apoptosis in cancer cells; promote angiogenesis and inflammation |
| | STC1 | Secreted by CAFs to enhance cancer cell intravasation and formation of distant metastases |
| | ISG15 | Regulate immune response upon stimulation by type I IFNs; may act on cancer cells to reinforce their invasive capacity and tumorigenic potential |
| | BCAT1 | Catalyse leucine transamination; may play a role in promoting cancer cell proliferation, migration and invasion |
| | OLFML2B | Potential oncogene related to regulation of the cell cycle, apoptosis and cell communication |

Supplementary Table. 2: Patient characteristics

| Age (years) | N=55(100%) |
|---|---|
| Median<br>Range | 51<br>27-72 |
| Sex | |
| Male<br>Female | 17 (30.9%)<br>38 (69.1%) |
| Tumour site | |
| Acral<br>Mucosal<br>Cutaneous<br>Unknown | 24 (43.6%)<br>8 (14.5%)<br>18 (32.7%)<br>5 (9.1%) |
| Tumour thickness | |
| ≤1 mm<br>>1-2 mm<br>>2-4 mm<br>>4 mm<br>Unknown | 0<br>4 (7.3%)<br>4 (7.3%)<br>18 (32.7%)<br>29 (52.7%) |
| Ulceration | |
| With<br>Without<br>Unknown | 22 (40.0%)<br>8 (14.5%)<br>25 (45.5%) |
| Metastasis status | |
| IIIC<br>M1a<br>M1b<br>M1c | 10 (18.2%)<br>16 (29.1%)<br>18 (32.7%)<br>11 (20.0%) |
| Efficacy | |
| CR<br>PR<br>SD<br>PD | 1 (1.8%)<br>13 (23.6%)<br>6 (10.9%)<br>35 (63.6%) |

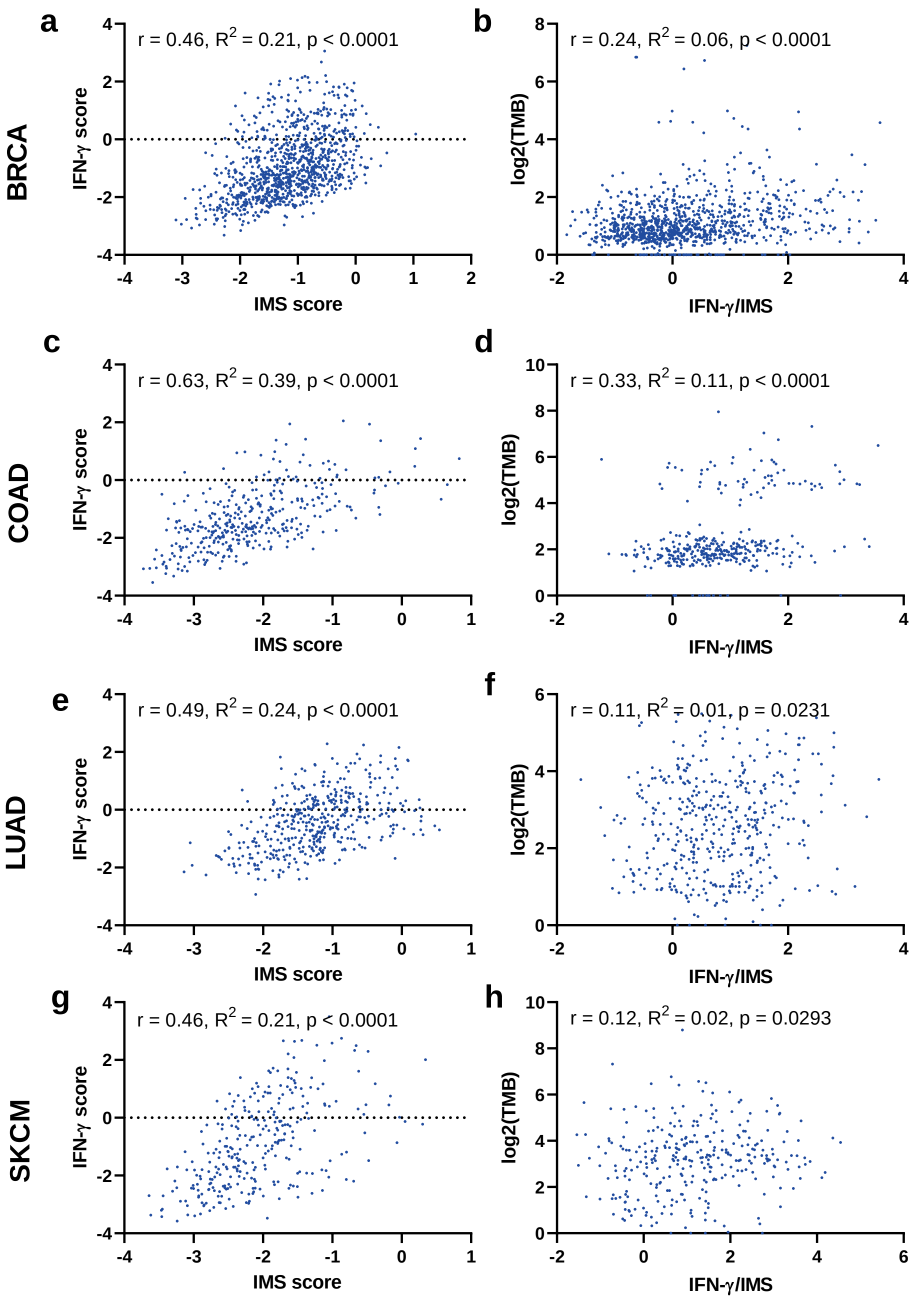


Supplementary Fig. 1:

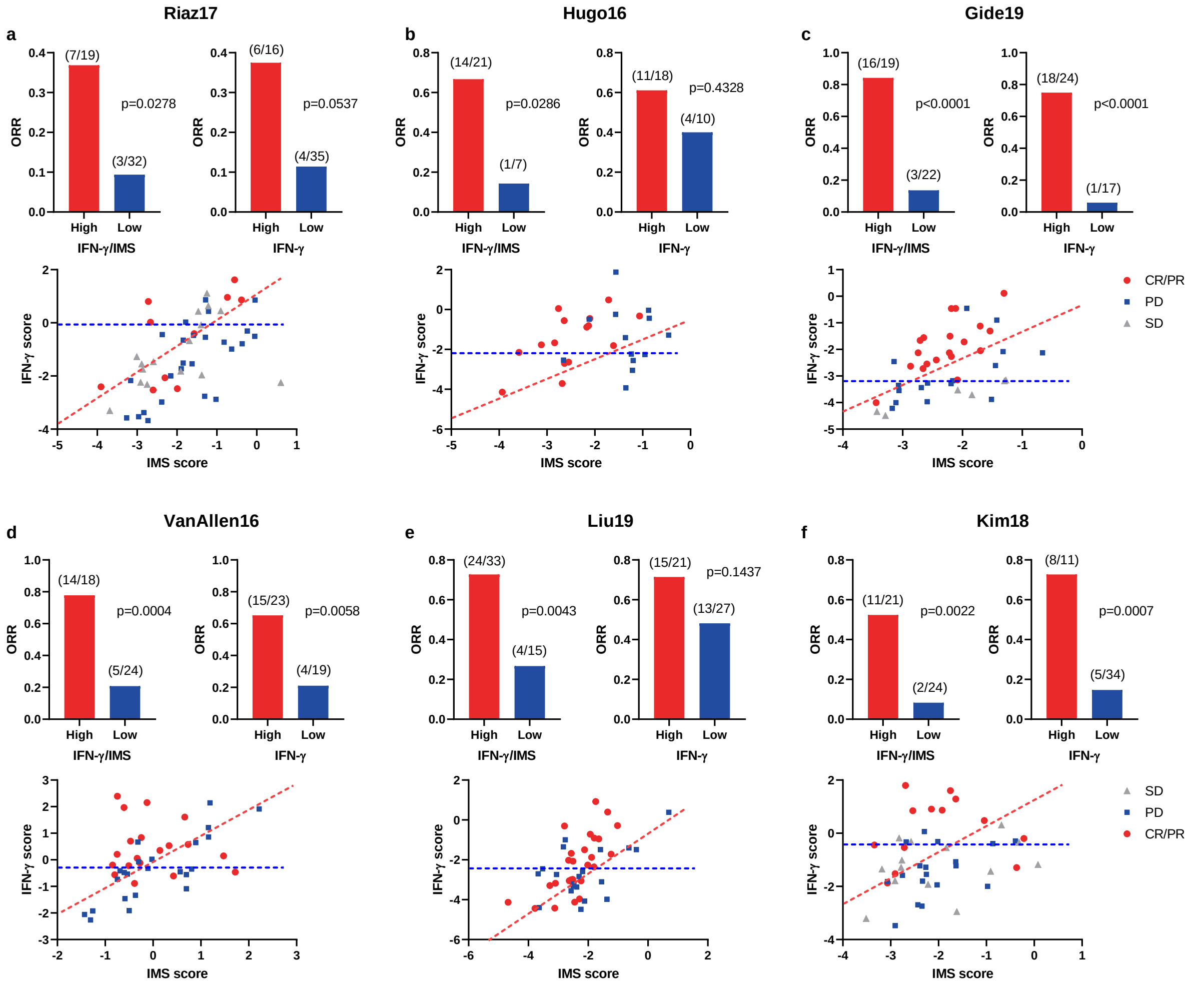


Supplementary Fig. 2:

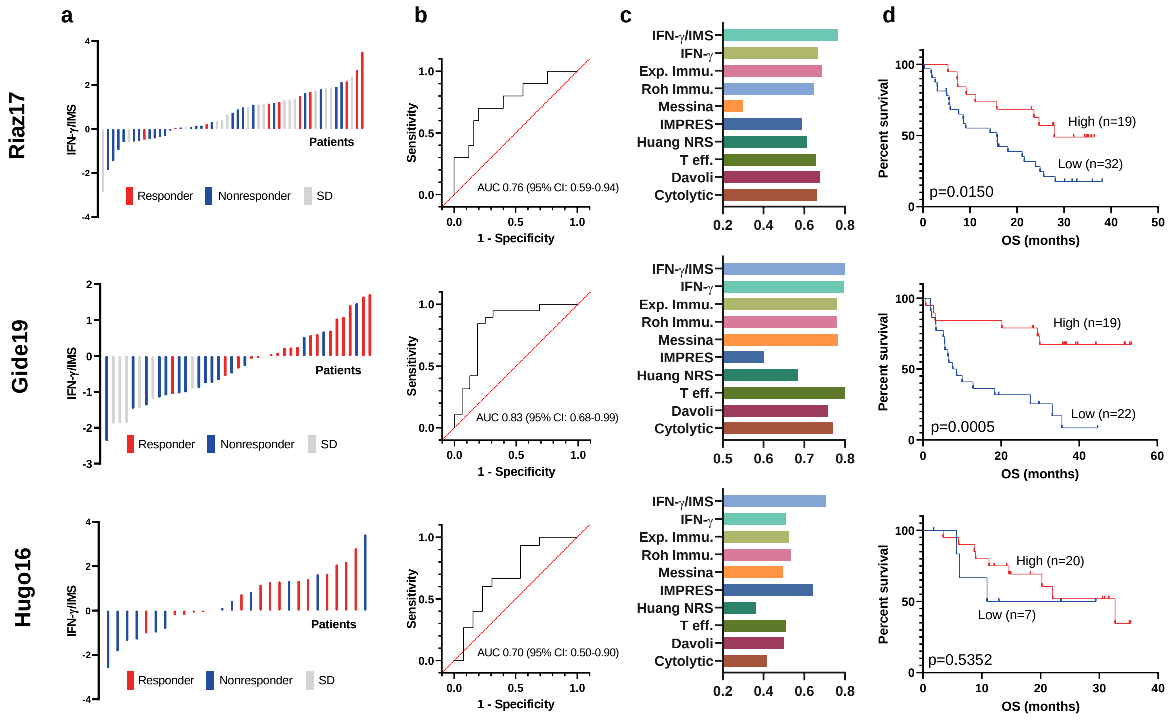


Supplementary Fig. 3:

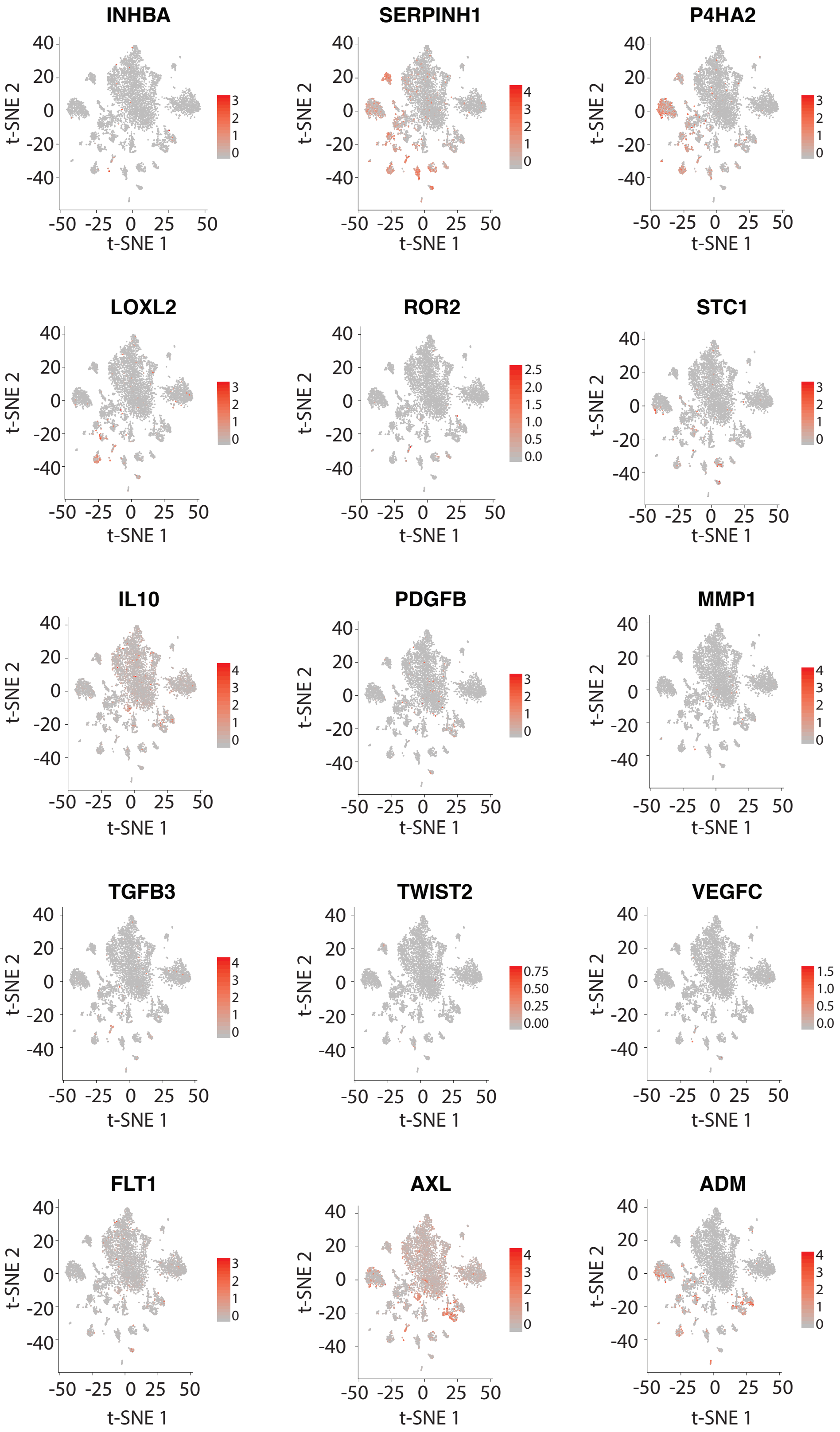


Supplementary Fig. 4:
